# Integrative Gene Co-expression Network Analysis Reveals Protein-Coding and lncRNA Genes Associated with Alzheimer’s Disease Pathology

**DOI:** 10.1101/2025.01.19.633755

**Authors:** Fatemeh Zebardast, Michael Peter Sascha Riethmüller, Katja Nowick

## Abstract

Alzheimer’s disease (AD) is a complex neurodegenerative disorder marked by widespread molecular changes, many of which remain poorly understood. While AD pathology progresses through specific brain regions, it is unclear whether these regions are affected similarly. Long non-coding RNAs (lncRNAs), emerging as key cellular regulators, remain largely uncharacterized in AD. Understanding how lncRNAs interact with protein-coding genes across brain regions could shed light on AD mechanisms and progression. To investigate this, we performed consensus weighted gene co-expression network analysis on 396 postmortem brain RNA-seq samples using a meta-analytic approach. Our analysis revealed substantial network rewiring in AD, particularly in the temporal cortex compared to the frontal cortex. The temporal cortex exhibited adaptive changes in gene interactions, while the frontal cortex showed a breakdown of healthy correlations—possibly reflecting regional differences in disease progression. We identified 46 protein-coding genes and 27 lncRNAs as key components in the AD network of the temporal cortex. Using known functions of protein-coding genes as reference points, we inferred potential functions for over 100 lncRNAs across both regions. These findings highlight novel lncRNA candidates potentially involved in AD and provide insights into their roles in both healthy and diseased brain states.

## Introduction

Alzheimer’s disease (AD) is a complex, multifactorial neurodegenerative disorder characterized by progressive cognitive decline and underlying pathological mechanisms (1). Current knowledge indicates that the hallmark pathology of AD begins in the medial temporal lobe (2) and, as the disease progresses, extends to other areas of the temporal cortex before spreading to additional brain regions, including the frontal lobes, in the mid to late stages (2–4). Its pathology disrupts numerous cellular mechanisms, such as mitochondrial functions (5–7), endoplasmic reticulum (8,9), Golgi apparatus integrity (10,11), cellular transport processes (5,12–14), and synaptic signaling (15–18). These widespread dysfunctions closely align with alterations in gene expression and molecular interactions, highlighting the complex biological networks that underlie AD.

Non-coding RNAs have attracted attention due to their potential involvement in both normal and disease-related cellular processes (19–22). Among these, long non-coding RNAs (lncRNAs)—transcripts longer than 200 nucleotides without apparent protein-coding capacity—have emerged as critical regulators of cellular functions (23–26). LncRNAs introduce an additional layer of complexity to gene regulatory processes through their interactions with protein-coding genes. Studies have implicated lncRNAs in several AD- associated dysfunctions, including amyloid beta (Aβ) production (27–30), oxidative stress (31), and synaptic impairments (32). Despite the annotation of numerous lncRNAs with known sequences and genomic locations, only a fraction have been extensively studied to uncover their functional roles in both healthy and disease conditions (24,26,33), leaving a significant knowledge gap yet to be addressed. A promising approach to address this gap is the construction of gene co-expression networks, which can reveal functional relationships between genes (34). Current methodologies face challenges such as mitigating noise and improving the reliability of gene-gene correlation detection (35,36). Overcoming these limitations is essential for generating accurate and interpretable biological insights. To address these challenges, we integrated gene co-expression networks derived from the same brain region across independent studies to construct consensus networks. This approach consolidates shared signals from multiple datasets, reducing the impact of dataset-specific noise and variability. By focusing on interactions consistently observed across networks, this method enhances the detection of biologically relevant signals, increasing confidence in their accuracy and interpretability compared to signals supported by individual networks alone (37–40).

In this study, we utilized RNA-seq data from two independent studies, comprising five datasets with 396 individuals total in AD and control groups, with gene expression profiles from the temporal cortex (TCX) and frontal cortex (FCX). These profiles were used to construct signed weighted topological overlap (wTO) networks, which were subsequently integrated into consensus networks to enhance reliability and biological relevance (Figure 1). Our analysis focused on lncRNAs and protein-coding genes, examining their correlations within gene co-expression networks under both AD and non-AD (control) conditions. Comparative analysis revealed significant rewiring of gene-gene interactions in the temporal cortex in AD, contrasted with a marked loss of ‘healthy’ gene correlations in the frontal cortex. Building on these observations, we identified 46 protein-coding genes and 27 lncRNAs as critical components of the AD network. A key objective of this study was to predict the functions of lncRNAs and their associations with AD pathology. By leveraging the established functions of protein-coding genes as network anchors, we assigned over 100 lncRNAs to biological processes in both AD and control conditions.

**Figure 1:**
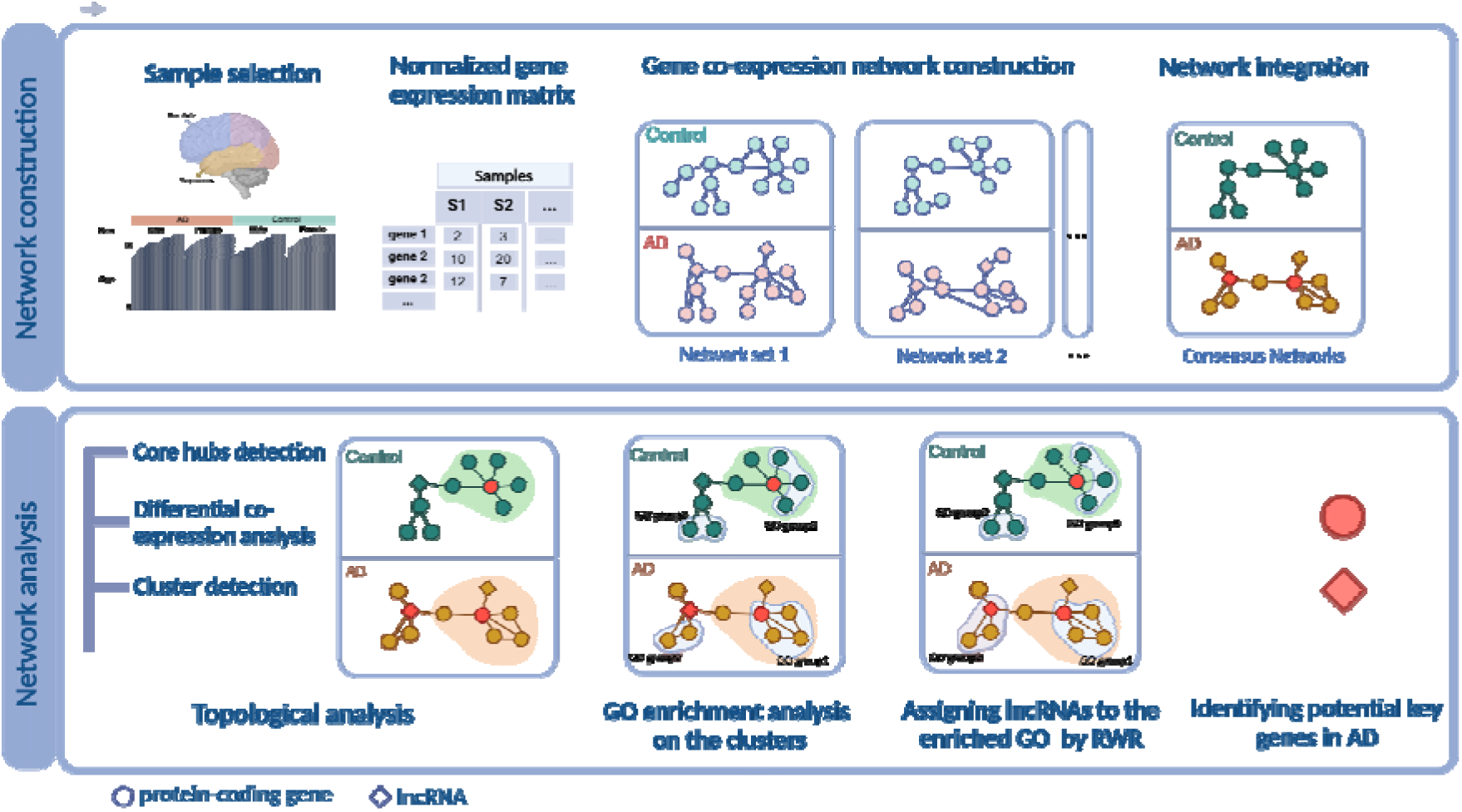
Overview and illustration of the current study. (41). From left to right, the top panel illustrates the network construction and integration process. The bottom panel illustrates the network analysis, including topology assessment, identification of hub genes, functional enrichment analysis, assigning enriched GOs to lncRNAs within each cluster, and identifying key genes associated with AD.

## Results

### Construction and integration of gene co-expression networks

We used a total of five RNA-seq datasets from two independent studies: the Mayo RNA-seq study (Mayo) (42) and the Mount Sinai Brain Bank study (MSBB) (43). The gene expression data were derived from two brain regions: TCX and FCX. These datasets were generated using samples with detailed clinical-pathological diagnoses, including Braak NFT stage (44), plaque density, CERAD (45), and CDR (46).

To minimize the impact of technical and unwanted biological factors, we implemented several steps: 1- Filtering individuals based on AD assessment criteria (Figure 2a). 2- Sex balance: each dataset includes an equal number of males and females. 3- Age balance: the age range was balanced within each group, as well as across all groups, as much as possible (Figure 2b-c). Furthermore, we uniformly processed all data from FASTQ files to minimize the impact of technical factors.

**Figure 2:**
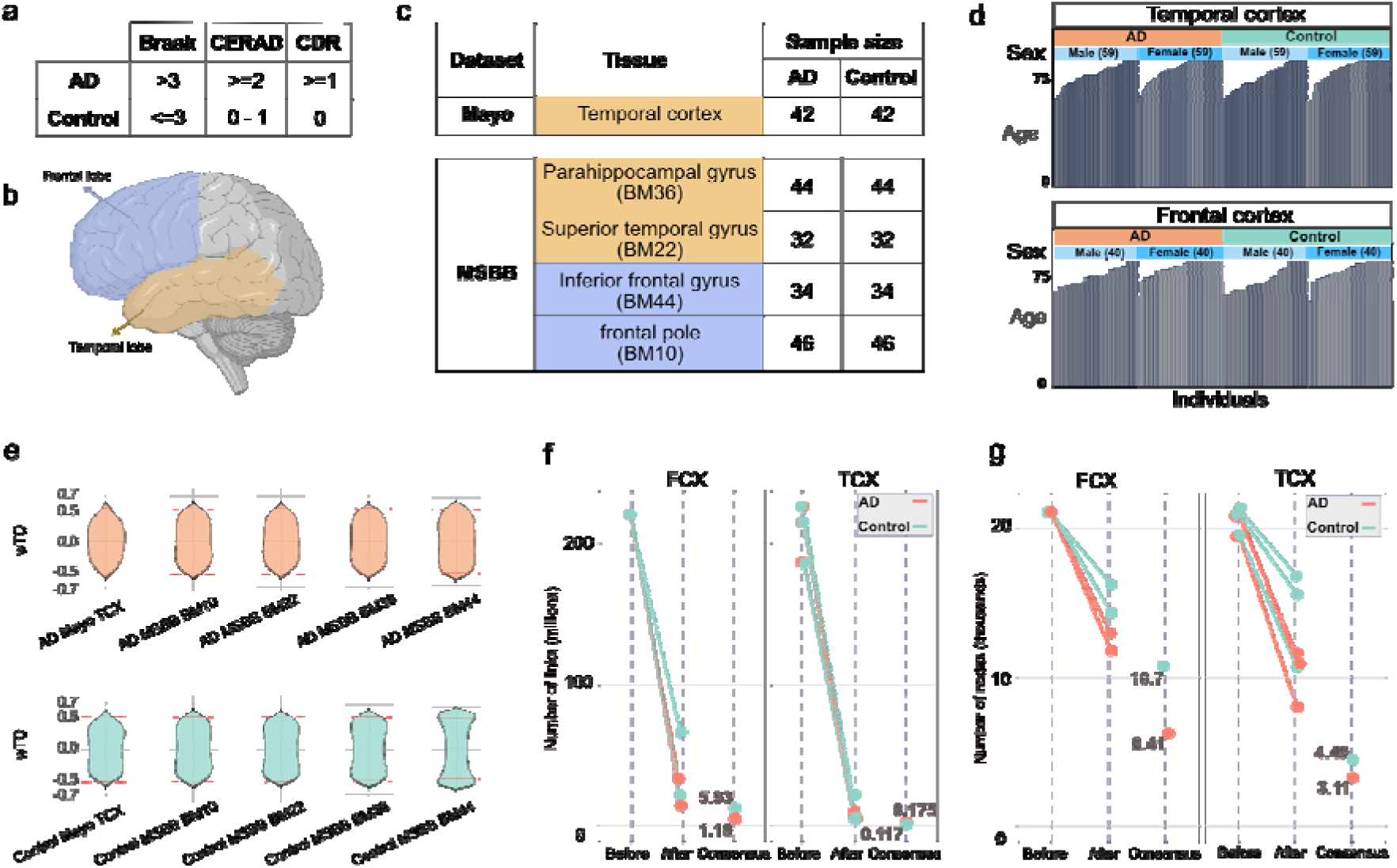
Summary of data demographics. **a)** Parameters and thresholds used to classify individuals into AD and control groups include Braak stage (based on neurofibrillary tangles distribution), CERAD (scoring neuritic plaques) from 0 (none) to 3 (frequent), and CDR (Clinical Dementia Rating) 0 (no dementia) to 3 (severe dementia). **b)** Illustration of brain areas used in this study that are temporal (yellow) and frontal (blue) cortices. **c)** The datasets, sample size, and Brodmann (BM) brain area utilized in this study. **d)** Data demography after data preparation: sample sizes were balanced between AD and control groups, as well as between sexes. Efforts were made to maintain a balanced age distribution. **e)** Distribution of wTO values for each network before filtering links for a cutoff. Red dashed lines indicate weight cutoffs. **f,g)** The graphs illustrate the number of links and nodes before and after filtering links and integration into consensus networks (x-axis: Before, After and Consensus). Each line and node represent one network, distinguished by colors for AD and control networks (see Tables S1 and S2 for more details).

Normalized gene count data were then used to construct signed weighted gene co-expression networks using the wTO method (47) separately for each AD and control dataset. We used gene count data for both protein-coding and lncRNA genes as input, resulting in networks that capture correlations between protein-coding and lncRNA genes. To focus on the strongest links, we considered the summary statistics of absolute weights of links in the networks (Figure 2e, supplementary Table S1) and excluded all weak links with an absolute weight below 0.5. This substantially reduced the number of links from a median of 219 million to 7.2 million links (Figure 2f). While the number of nodes also decreased, the reduction was less drastic, dropping from 20,900 nodes before filtering to a median of 13,200 nodes afterward (Figure 2g). This implied that most genes remained in the network, but only the most robust interactions were represented.

Subsequently, the filtered networks were integrated separately for AD and control groups based on brain regions to further enhance the reliability of the networks, excluding links that might be dataset/study specific. Specifically, the Mayo TCX network was integrated with two MSBB networks, BM36 and BM22, all derived from different parts of the temporal cortex, into a consensus TCX network. Similarly, the BM10 and BM44 networks, which originated from two distinct areas of the frontal cortex, were integrated to form the consensus FCX network (Figure 2 b-c, supplementary Table S2). The resulting consensus networks included only the commonalities across the integrated networks for each brain region. This method ensured that only gene relations observed in multiple studies were included in our further analyses. From here on, we refer to these consensus networks as the “AD” and “control” networks of the TCX and FCX, respectively.

### Comparative analysis of the TCX networks

We began our analysis with the TCX networks. Using K-core decomposition, we identified nodes in the innermost shell as core genes (hubs) in both AD and control networks (supplementary Table S3 and S4). The TCX AD network had 306 core genes, compared to 235 in the TCX control network, with 18% (81 genes) shared between them (Figure 3a). Notably, lncRNAs accounted for 15% of the core genes in the AD network, nearly double the 8% observed in the control network (Figure 3b). Statistical evaluation using Fisher’s exact test confirmed the significant enrichment of core lncRNAs in the AD network (p-value = 0.016, odds ratio = 2.01). Among the TCX AD core genes, 22 were significantly differentially expressed (DE, FDR < 0.05 and absolute log fold change> 0.3) across all three TCX datasets, all of which were downregulated in AD (Figure 3 c and d). These included 17 protein-coding and 5 lncRNA core genes. A complete list of DE genes is provided in supplementary Table S5.

**Figure 3:**
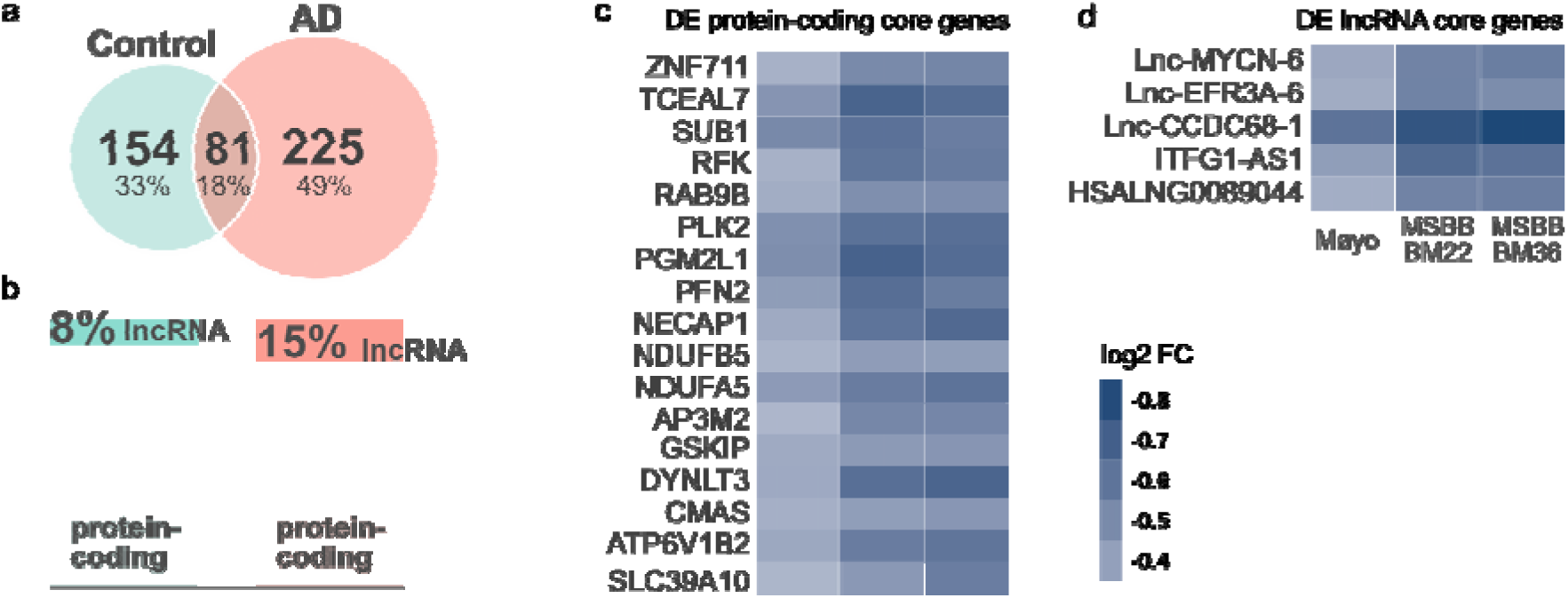
Demography and differential expression analysis of TCX core genes. **a)** Venn diagram depicting the overlap of core genes between AD (orange) and control (green) networks in TCX. **b)** Proportion of protein-coding and lncRNA genes within cores of TCX AD network (orange) and control network (green). **c** and **d)** Heatmaps showing the log2 fold change (FC) for differentially expressed (DE) core genes in AD, including protein-coding (c) and lncRNA (d) genes that are in the TCX AD network. Each column represents the log2 FC for a single dataset used to construct the TCX networks, which are Mayo, MSBB BM22 and BM36.

To examine the structure and underlying biological functions of the networks, we applied the Louvain method for community detection to each consensus network separately. In the TCX AD network, we observed three gene clusters, whereas the control network formed five clusters (Figure 4a, Table 1). GO enrichment analysis was conducted on the protein-coding genes within each cluster, using the same set of background genes (see Methods). Enriched GO terms were manually curated and grouped into broader functional categories to address GO term redundancy (see Methods for details). Significantly enriched biological processes fell into eight broad functional categories, including 1- mitochondria energy production, 2- intracellular trafficking, 3- synapse signaling, 4- neurogenesis, 5- apoptosis, 6- cytoskeleton organization 7- gene expression, 8- cellular homeostasis, which were enriched to differing degrees among the clusters (Figure 4b, supplementary Table S6). Additionally, four GO terms were grouped as “others” due to their specificity or broad relevance (e.g., GTPase activity). Among all clusters, only genes of control cluster 5 were not significantly enriched for any terms. Cellular homeostasis was the only functional category that was enriched only in control clusters.

**Figure 4:**
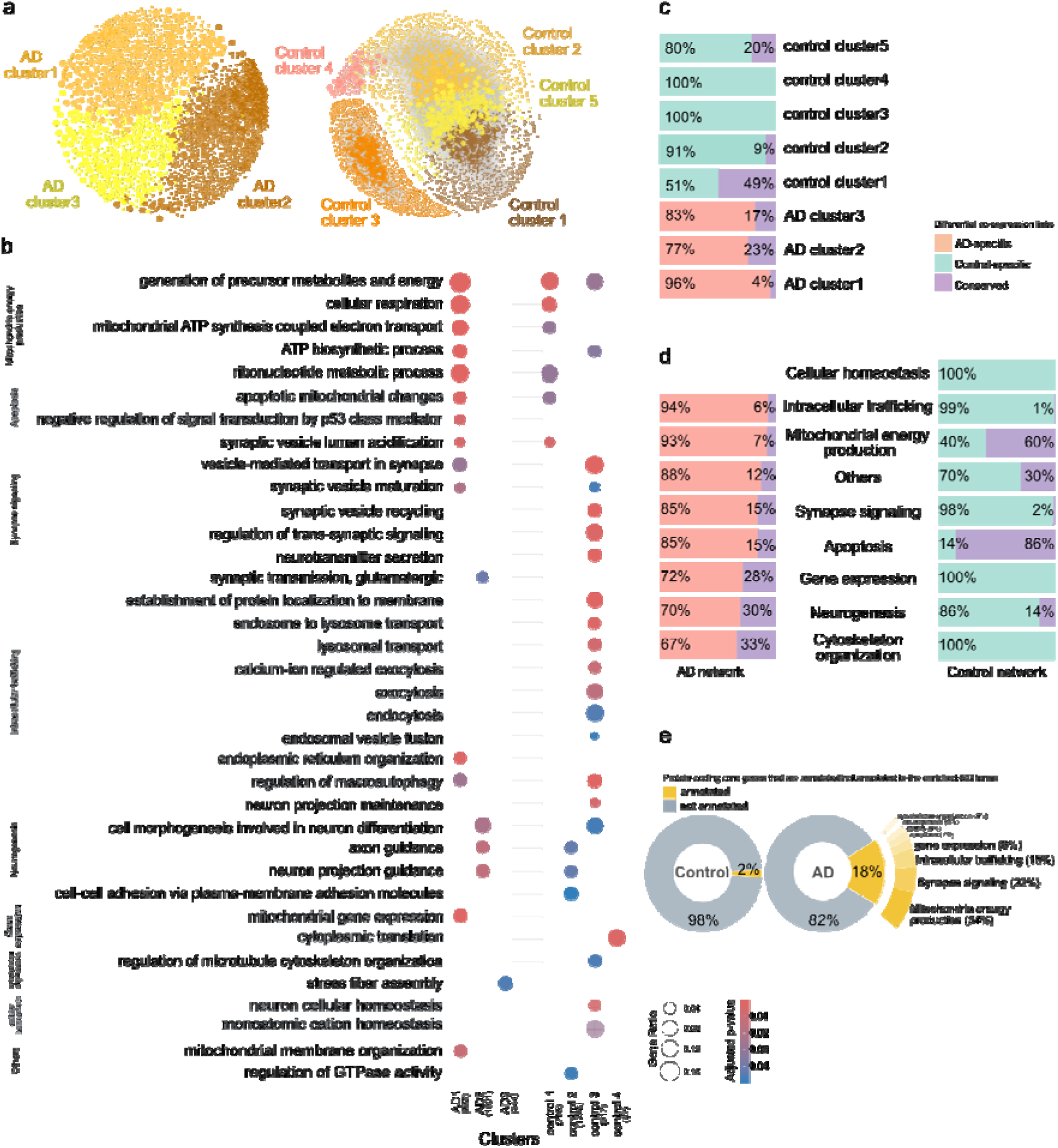
Topological and differential co-expression analysis of TCX networks. **a)** Representation of clusters for TCX AD and control networks, shown on the right and left, respectively. There are two node sizes; larger nodes represent core hub genes, while smaller nodes represent the remaining genes. Each color represents a distinct cluster. Gray backgrounds indicate links, which are faded to reduce complexity for visualization. The networks were visualized by Gephi (48). **b)** The plot displays selected enriched GO terms and their classification into broader functional categories. The significance of enriched GOs was assessed using the false discovery rate, and terms with an adjusted p-value below 0.05 were selected. Dot colors indicate the adjusted p-value, and dot size is proportional to the number of genes of interest in the input gene list for the given term. **c)** Distribution of differential co-expression link categories within each cluster. The percentages represent relative abundance, meaning the original proportion of link types within each cluster is normalized relative to the cluster size. **d)** Distribution of differential co-expression links categories connecting genes associated with each functional category. **e)** Distribution of protein-coding core genes annotated to the enriched GO terms.

**Table 1:** The topological details of AD and control clusters in the temporal cortex.

|  | Cluster 1 |  | Cluster 2 |  | Cluster 3 |  | Cluster 4 |  | Cluster 5 |  |
| --- | --- | --- | --- | --- | --- | --- | --- | --- | --- | --- |
|  | Nodes | Links | Nodes | Links | Nodes | Links | Nodes | Links | Nodes | Links |
| **control** | 961 | 18,583 | 1,662 | 31,509 | 836 | 4,702 | 113 | 169 | 894 | 30,439 |
| **AD** | 920 | 48,198 | 1,689 | 44,374 | 476 | 17,262 |  |  |  |  |

In order to gain a deeper understanding of the differences between AD and control networks, a differential co-expression network analysis was performed (49). We classified links into three categories: 1- conserved links, which were common between the two conditions (i.e., AD and control); 2- condition-specific links, which appeared only in one condition (i.e., AD-specific); and 3- reversed links, which represented correlations with opposite signs in the two conditions. The results showed that 96%, 77%, and 83% of links in AD clusters 1, 2, and 3, respectively, were AD-specific (Figure 4c). This pattern was also reflected in links between genes of each functional category in AD clusters (Figure 4d). These findings indicated that AD-related changes are not confined to a single cluster or GO term; rather, they broadly impacted all three gene clusters and the identified underlying biological process. Similar observations were seen in the control network analysis, indicating that the majority of links per cluster and functional category were control-specific and, in other words, were absent in AD. There was only one exception: 51% of control cluster 1 included conserved links, which involved genes annotated to mitochondria energy production and apoptosis.

The observed rewiring of the AD network suggested a shift in the organization of molecular interactions. To investigate the key players of this rewiring, we examined all core genes, as these genes were likely central to the disrupted processes and extensive rewiring observed in the AD network. Notably, of the protein-coding core genes, a higher proportion was annotated to the functional categories in the AD network (18%, 46 genes) compared to the control network (2%, 5 genes) (Figure 4e). Among the core genes linked to the enriched GOs, the largest proportion (34%) were associated with mitochondrial energy production, followed by 22% with synaptic signaling and 15% with intracellular trafficking (Figure 4e). While the majority of the protein-coding core genes in both networks were not annotated to enriched GO terms, the increased representation of AD core genes in these categories underscored their potential roles in key processes disrupted in AD pathology, particularly in energy production, neural communication, and intracellular transport.

Among the 46 core genes of the AD network annotated to the enriched GO terms (Table 2, Figure 5), nearly all were exclusively core to the AD network, except for four genes. Notably, these four genes—*CHL1* and *USP33* (neurogenesis), *GRIA2* (synapse signaling), and *YARS2* (mitochondrial gene expression)—were annotated to biological processes only within the AD network and not in the control network. This observation suggests that these genes might gain a disease-related role in the context of AD. It might be observed that a single gene was annotated to different functional categories in the AD and control networks (Table 2). This was explained by the fact that the gene clustered with different sets of genes in the two networks, leading to its association with distinct GO terms. This observation reflected the differing patterns of gene interactions between AD and control networks, likely linked to the disease condition.

**Figure 5:**
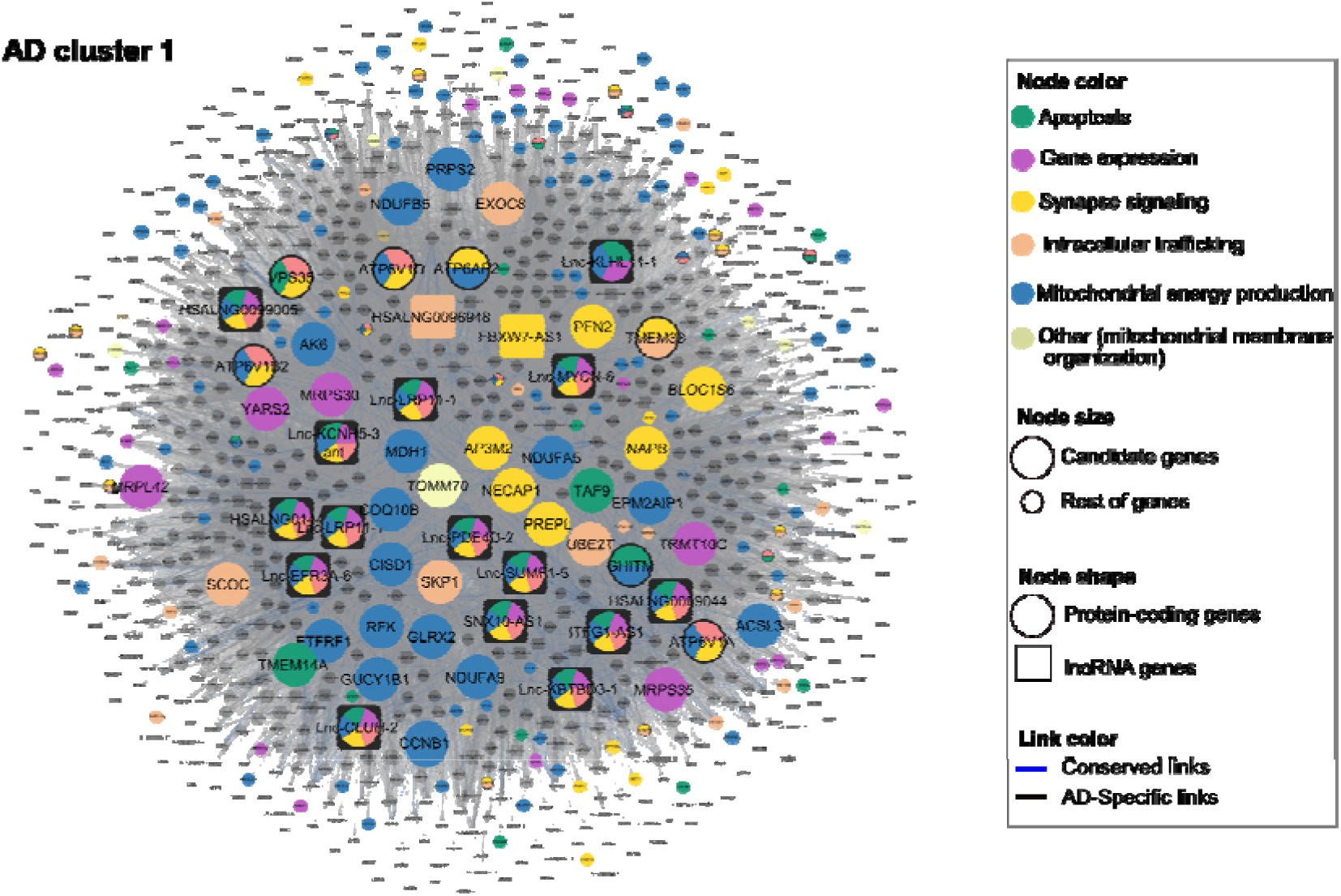
AD cluster 1 network visualization. Node colors indicate functional categories; multicolored nodes represent genes annotated to multiple categories. Larger nodes are identified as key genes in the AD network, while smaller nodes represent other genes in AD cluster 1. Circles represent protein-coding genes, and squares represent lncRNA genes. Link colors: blue links are conserved between the TCX AD and control network, and black links are AD-specific links. Names of key genes are enlarged; names of other genes are visible upon full zoom. Further details can be explored through the interactive Shiny app we have provided.

**Table 2:** Identified key protein-coding genes potentially associated with AD pathology in TCX.

| Gene name | KEGG pathway | DEG | AD |  | Control |  |  |
| --- | --- | --- | --- | --- | --- | --- | --- |
|  |  |  | Cluster | Functional category | Cluster | Functional category | hub |
| *ABL1* | - | - | 3 | Cytoskeleton organization | 1 | - | - |
| *ACSL3* | - | - | 1 | Mitochondrial energy production | 1 | Mitochondrial energy production | - |
| *AK6* | - | - | 1 | Mitochondrial energy production | 1 | Mitochondrial energy production | - |
| *AP3M2* | - | Down | 1 | Synapse signaling | 1 | - | - |
| *ATL1* | - | - | 2 | Neurogenesis | - | - | - |
| *ATP6AP2* | - | - | 1 | Mitochondrial energy production, synapse signaling | 1 | Intracellular trafficking, synapse signaling | - |
| *ATP6V1A* | - | - | 1 | Intracellular trafficking, mitochondrial energy production, synapse signaling | 1 | Intracellular trafficking, mitochondrial energy production, synapse signaling | - |
| *ATP6V1B2* | - | Down | 1 | Intracellular trafficking, mitochondrial energy production, synapse signaling | 1 | Intracellular trafficking, Mitochondrial energy production, synapse signaling | - |
| *ATP6V1D* | - | - | 1 | Intracellular trafficking, mitochondrial energy production, synapse signaling | 1 | Intracellular trafficking, synapse signaling | - |
| *BLOC1S6* | - | - | 1 | Synapse signaling | 1 | - | - |
| *CCNB1* | - | - | 1 | Mitochondrial energy production | 1 | Mitochondrial energy production | - |
| *CHL1* | - | - | 2 | Neurogenesis | 5 | - | core |
| *CISD1* | - | - | 1 | Mitochondrial energy production | 1 | Mitochondrial energy production | - |
| *COQ10B* | - | - | 1 | Mitochondrial energy production | 1 | Mitochondrial energy production | - |
| *EPM2AIP1* | - | - | 1 | Mitochondrial energy production | 1 | Mitochondrial energy production | - |
| *ETFRF1* | - | - | 1 | Mitochondrial energy production | 1 | Mitochondrial energy production | - |
| *EXOC8* | - | - | 1 | Intracellular trafficking | 1 | - | - |
| *GHITM* | - |  | 1 | Apoptosis, mitochondrial energy production, others (mitochondrial membrane organization) | 1 | Apoptosis, mitochondrial energy production | - |
| *GLRX2* | - |  | 1 | Mitochondrial energy production | 1 | Mitochondrial energy production | - |
| *GRIA2* | - |  | 2 | Synapse signaling | 2 | - | core |
| *GUCY1B1* | - |  | 1 | Mitochondrial energy production | 1 | Mitochondrial energy production | - |
| *MDH1* | - |  | 1 | Mitochondrial energy production | 1 | Mitochondrial energy production | - |
| *MKKS* | - |  | 3 | Cytoskeleton organization | 1 | - | - |
| *MRPL42* | - |  | 1 | Gene expression | 1 | - | - |
| *MRPS30* | - | - | 1 | Gene expression | 1 | - | - |
| *MRPS35* | - | - | 1 | Gene expression | 1 | - | - |
| *NAPB* | - | - | 1 | Synapse signaling | 1 | - | - |
| *NDUFA5* | AD pathway | Down | 1 | Mitochondrial energy production | 1 | Mitochondrial energy production | - |
| *NDUFA9* | AD pathway | - | 1 | Mitochondrial energy production | 1 | Mitochondrial energy production | - |
| *NDUFB5* | AD pathway | Down | 1 | Mitochondrial energy production | 1 | Mitochondrial energy production | - |
| *NECAP1* | - | Down | 1 | Synapse signaling | 1 | - | - |
| *PFN2* | - | Down | 1 | Synapse signaling | 1 | - | - |
| *PREPL* | - | - | 1 | Synapse signaling | 1 | - | - |
| *PRPS2* | - | - | 1 | Mitochondrial energy production | 1 | Mitochondrial energy production | - |
| *RFK* | - | Down | 1 | Mitochondrial energy production | 1 | Mitochondrial energy production | - |
| *SCOC* | - | - | 1 | Intracellular trafficking | 1 | - | - |
| *SKP1* | - | - | 1 | Intracellular trafficking | 1 | - | - |
| *TAF9* | - | - | 1 | Apoptosis | 1 | - | - |
| *TMEM14A* | - | - | 1 | Apoptosis, others (mitochondrial membrane organization) | 1 | Apoptosis | - |
| *TMEM33* | - | - | 1 | Intracellular trafficking, synapse signaling | 1 | - | - |
| *TOMM70* | - | - | 1 | others (mitochondrial membrane organization) | 1 | - | - |
| *TRMT10C* | - | - | 1 | Gene expression | 1 | - | - |
| *UBE2T* | - | - | 1 | Intracellular trafficking | 1 | - | - |
| *USP33* | - | - | 2 | Neurogenesis | 5 | - | core |
| *VPS35* | - | - | 1 | Apoptosis, intracellular trafficking, synapse signaling | 1 | Apoptosis | - |
| *YARS2* | - | - | 1 | Gene expression | 1 | - | core |

The table lists 46 protein-coding key genes potentially associated with AD pathology in the TCX AD network. Various features of these genes are listed, including their involvement in the KEGG AD pathway, differential expressions in AD (DEG), cluster membership, as well as their annotation to the functional categories. All are AD hubs, while their hub status in control is indicated under the hub column.

### lncRNA functional assignment and core lncRNAs in TCX networks

The gene co-expression networks in this study included both protein-coding and lncRNA genes, providing us with the opportunity to gather functional information for lncRNAs based on their network neighbors. For each functional category, Random Walk with Restart (RWR) was run using annotated protein-coding genes as seeds. LncRNAs were ranked based on their RWR scores, and the top 10th percentile were assigned to the corresponding functional category. To ensure the biological relevance of these assignments, a leave-one-out approach was applied to validate the rankings. A lncRNA was assigned to a category only if its RWR score exceeded the lowest score from the leave-one-out simulation (Methods, supplementary Figure S1 and Table S8). Using this approach, 54 lncRNAs in the TCX AD network and 103 in the TCX control network were assigned to functional categories within their respective clusters (supplementary Table S8).

Out of the 46 AD core lncRNA genes, we focused on 27 lncRNAs that were assigned to at least one functional category, as these lncRNAs likely played key roles in the dysregulation of their assigned functional categories within the AD network (Table 3, Figure 5). All but six of these lncRNAs were AD-specific core genes. Notably, *FBXW7-AS1*, *Lnc-PDE4D-2*, *Lnc- KLHL11-1*, *Lnc-NIF3L1-5*, *Lnc-KALRN-1*, *Lnc-CCDC68-1*, *HSALNG0099005*, *HSALNG0105911*, *ZRANB2-DT*, *ENSG00000287022*, and *SLC8A1-AS1* were functionally assigned only within the AD network, suggesting a gain of function for these lncRNAs in the context of AD. The remaining lncRNAs were functionally assigned in both networks, either to similar or different functional categories (Table 3). Many of these lncRNAs were assigned to all functional categories within their respective clusters, as expected given their status as core genes with high connectivity to numerous nodes, which often results in high RWR scores. However, a closer examination of their functional categories and comparisons with the control network revealed considerable differences. This suggests that lncRNAs were incorporated into different neighborhoods or changed the sets of genes they regulated in AD compared to control brains. For example, *POC1B-AS1* was assigned to cytoskeleton organization-related processes in the AD network, whereas it was associated with mitochondrial energy production in the control network. Among these 27 core lncRNAs, five were downregulated in AD, namely *ITFG1-AS1*, *Lnc-MYCN-6*, *Lnc-EFR3A-6*, *HSALNG0089044*, and *Lnc-CCDC68-1* (Figure 3d, Table 3).

**Table 3:**
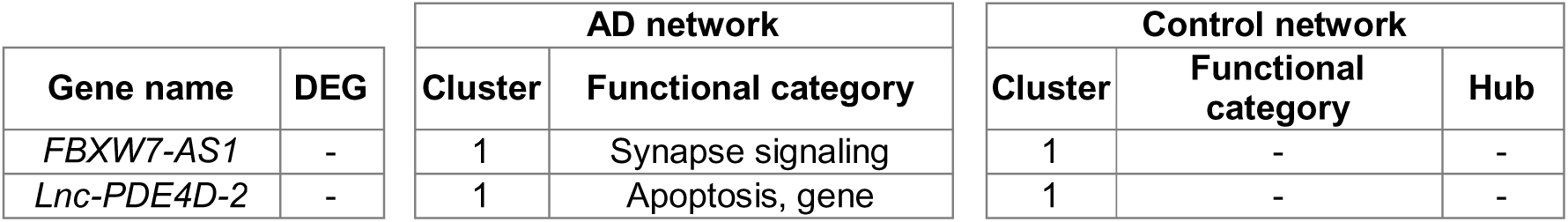
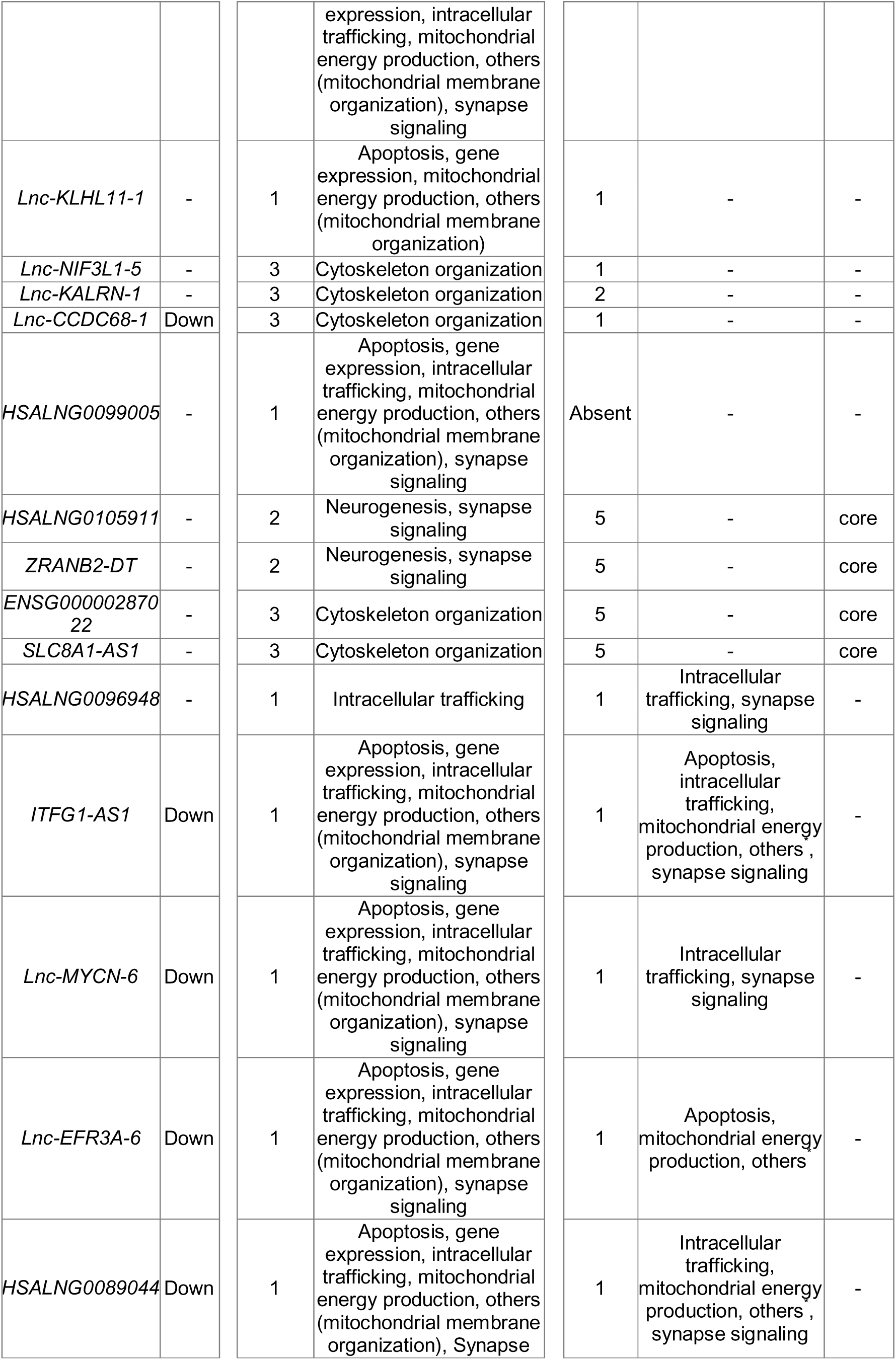
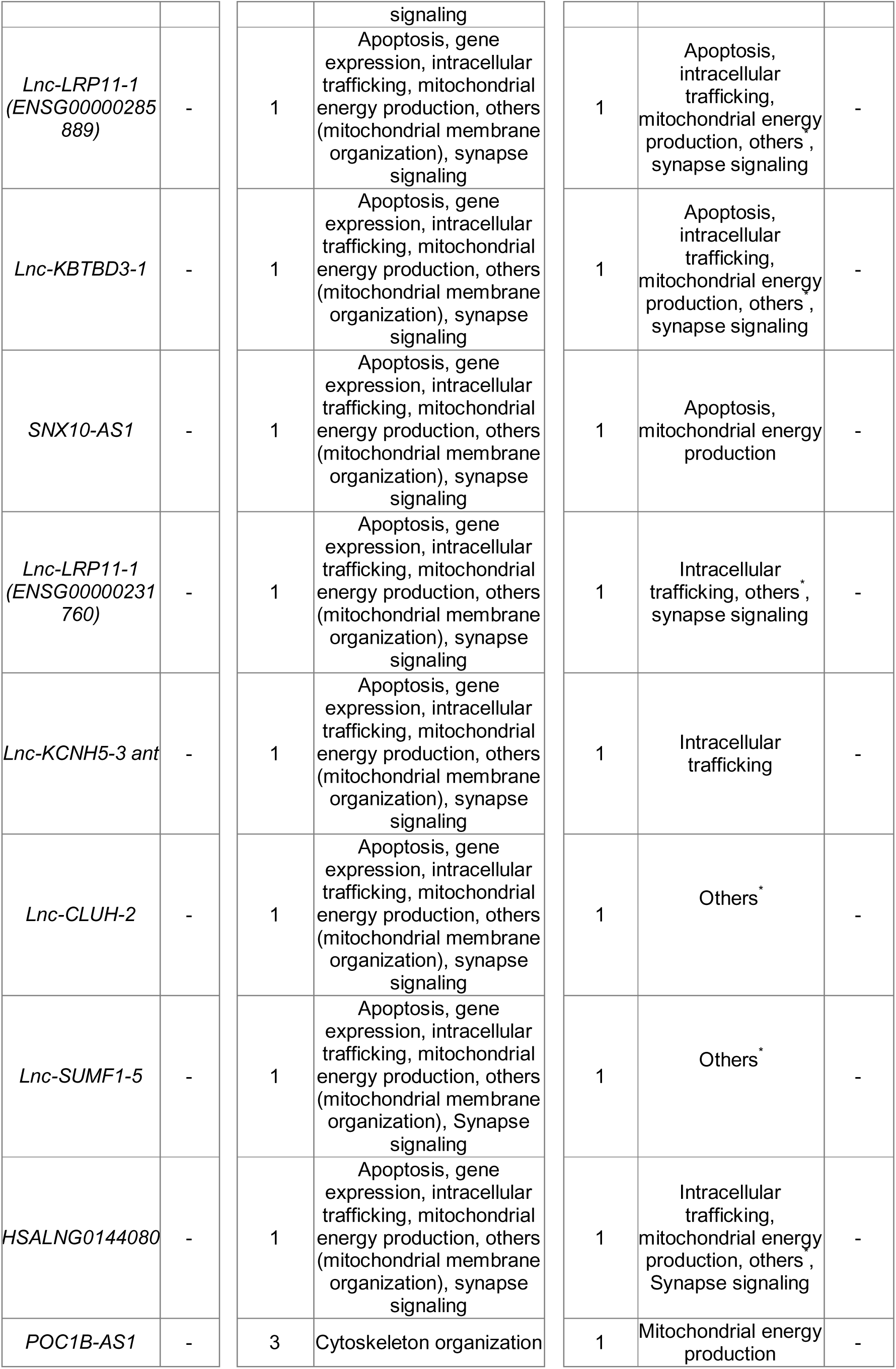
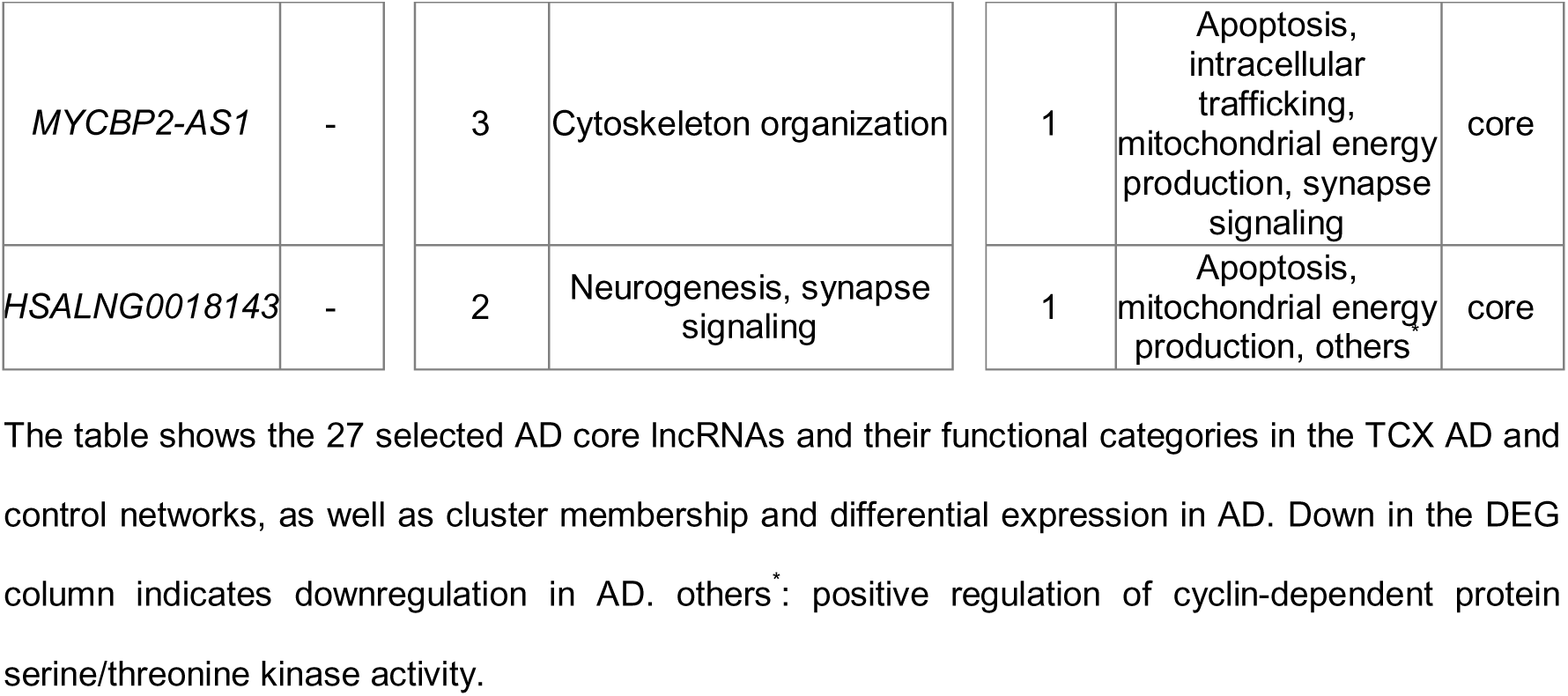
Key lncRNA genes potentially associated with dysregulated GO terms in the TCX AD network.

### Comparing TCX and FCX

After analyzing networks of the temporal cortex, we performed the same analysis for the frontal cortex and compared our results between the two brain areas. In the frontal cortex, we found that unlike in the temporal cortex, the AD network shows much higher similarity to the control network (Figure 6 a-d), where the majority of links are conserved in the AD network (86%, figure 6a). Conversely, the control network is predominantly composed of control-specific links (83%), with minimal overlap (14%) with links found in the AD network (Figure 6b). This observation implied that the AD network in the frontal cortex has lost a significant number of gene-gene correlations. This hypothesis was supported by the marked difference in the size of the networks: the AD network contained 1.17 million links, while the control network contained 5.93 million links. This pattern was also evident in the significant overlap between AD core hubs and control core genes (Figure 6e). While this overlap suggested that certain gene relationships were preserved in AD, despite the extensive loss of specific gene-gene interactions, a substantial number of core genes presented in the control network were absent in the AD network. Interestingly, the proportion of gene types, i.e., protein-coding or lncRNA genes, among the core genes {Citation}was very similar between AD and control networks, with 15% or 18% of cores being lncRNAs, respectively (Figure 6f).

**Figure 6:**
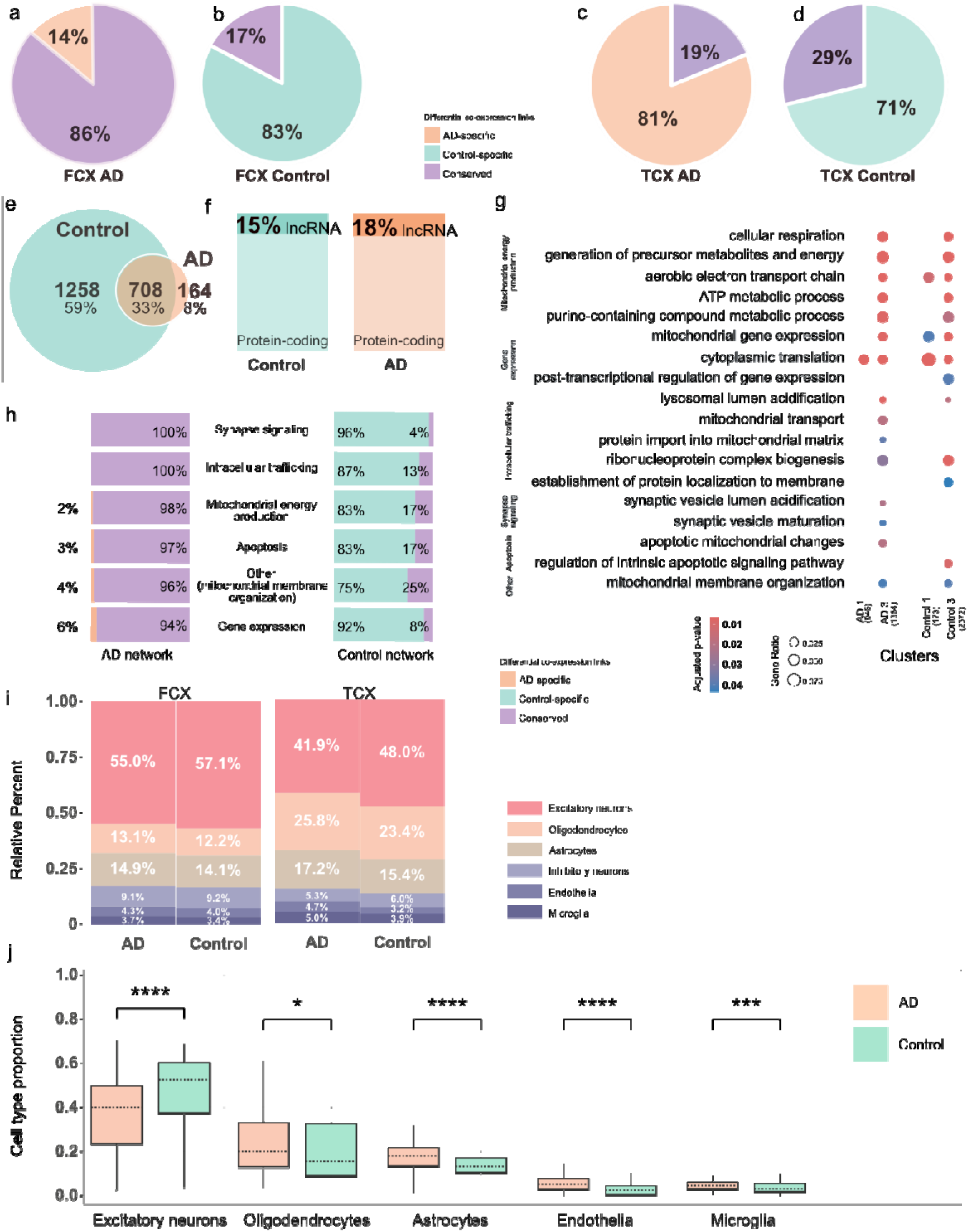
Results of comparative analysis between FCX and TCX. Pie chart showing proportions of differential co-expressed links **a)** in the FCX AD network, **b)** FCX control network, **c)** TCX AD network and **d)** TCX control network. **e** Venn diagram showing the overlap of core genes between AD and control networks in FCX. **f)** Proportion of protein-coding and lncRNA genes within FCX core genes. **g)** Significantly enriched GO terms for biological processes, grouped into five broader functional categories: 1) mitochondrial energy production, 2) intracellular trafficking, 3) synapse signaling, 4) apoptosis, 5) gene expression, with an additional “other” category with a single term-mitochondrial membrane organization. **h)** Distribution of differential link categories within each functional category, with percentages representing relative abundance normalized to cluster size. **i)** Stacked bar plots showing the estimated cell-type proportions in FCX and TCX for AD and control samples. Each color represents one cell type. **j)** Significant cell-type proportion changes between AD and controls were observed only in TCX datasets. P-values were measured by the Wilcoxon test (*p < 0.05, **p < 0.01, ***p < 0.001, ****p < 0.0001).

We used the same method as for the TCX to cluster the FCX consensus networks (supplementary Table S9). As done for the TCX, significantly enriched GO terms were then manually classified into five broader functional categories (Figure 6g) including: 1- mitochondrial energy production, 2-intracellular trafficking, 3- synapse signaling, 4- apoptosis, 5- gene expression. Additionally, an “other” category was included, containing a single term, mitochondrial membrane organization. A complete list of enriched GO terms can be found in supplementary table S10. Genes in clusters 1 and 3 of both AD and control networks were significantly enriched for biological process GO terms, but not in the other clusters.

Differential gene co-expression analysis revealed a dominance of conserved links connecting genes associated with enriched GO terms within AD clusters (Figure 6h). Each functional category showed 94% to 100% conserved links. In contrast, in the control network, the majority of links were control-specific within the functional categories. One possible interpretation of this tendency was that the pathogenic alterations associated with AD disrupted numerous healthy gene expression correlations in the frontal cortex area. This might also suggest that AD-specific correlations represent cellular compensation against AD-related disruptions.

In bulk RNA-Seq data, observed differences in gene expression can often be attributed to variations in cell-type composition. Given that such differences might also contribute to alterations in gene co-expression networks, we sought to assess the cellular composition of the bulk RNA-seq data used in this study. To this end, we performed a deconvolution analysis on both the TCX and FCX regions using CIBERSORT, utilizing human brain single-cell data as a reference (Figure 6i). Across both brain regions and independent of disease status, excitatory neurons were the predominant cell type, comprising 41–56% of the total population. However, the relative abundance of other cell types varied between TCX and FCX, with microglia consistently being the least abundant in both regions. The overrepresentation of excitatory neurons and the underrepresentation of microglia align with previous findings in the human brain, indicating that bulk RNA-seq data primarily capture transcriptional changes in excitatory neurons, while microglia, due to their lower RNA content, are typically underrepresented (50).

To assess whether AD pathology affects cell-type composition, we compared cell-type proportions between AD and control samples within each brain region. Interestingly, we observed significant alterations only in TCX, where excitatory neurons were markedly decreased (p.adj = 9.0 × 10⁻LJ), while astrocytes (p.adj = 4.66 × 10⁻LJ), microglia (p.adj = 5.0 × 10⁻LJ), oligodendrocytes (p.adj = 0.03), and endothelial cells (p.adj = 2.07 × 10⁻LJ) were significantly increased (Figure 6j). This shift in cellular composition may contribute to the gene co-expression rewiring observed in the TCX AD network.

In addition, the FCX control network was employed to evaluate the reliability and consistency of the lncRNA annotation method implemented in this study. We investigated the similarity of lncRNAs annotated to the same functional category between the TCX and FCX control networks (supplementary Table S11). Five functional categories common to both networks were analyzed, including mitochondrial energy production, intracellular trafficking, synapse signaling, apoptosis, and gene expression (supplementary Table S12). Significant overlaps of lncRNAs assigned to functional categories were observed between the FCX and TCX control networks, with p-values approaching zero (Table 4). The analysis accounted for the disparity in lncRNA totals between FCX and TCX control networks (1100 and 504, respectively) and a permutation test for statistical significance. Notably, the gene expression category was the only one without any overlapping lncRNAs. Additionally, permutation tests were conducted for lncRNAs not assigned to functional categories. These tests yielded a non-significant p-value of 0.71, highlighting that the functional assignments of annotated lncRNAs are significantly more consistent than would be expected by chance.

**Table 4:**
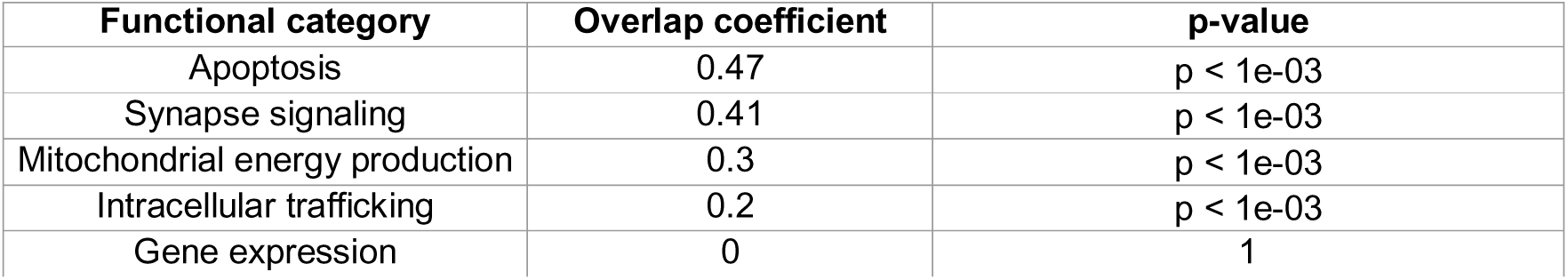
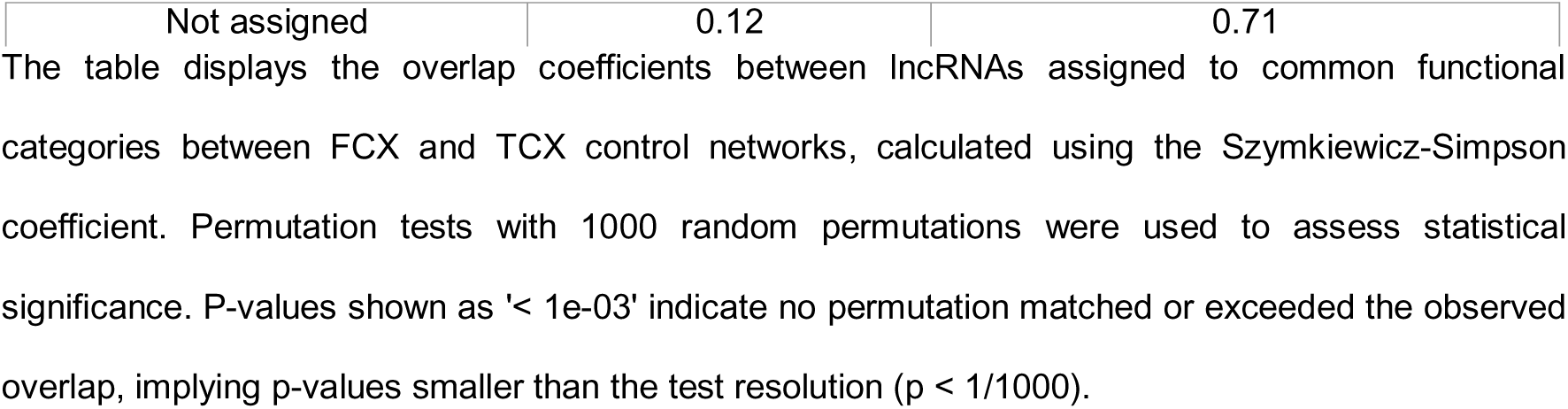
Similarity of lncRNA annotations across FCX and TCX control networks.

## Discussion

In this study, we presented robust consensus gene co-expression networks including protein-coding and lncRNA genes of two brain regions (temporal cortex and frontal cortex) and investigated how the networks differ in AD compared to control brains. Our deep comparative analysis between the consensus networks of AD and control groups highlighted significant rewiring of gene-gene interactions within the temporal cortex area (Figure 4c, d), in contrast to a more pronounced loss of ‘healthy’ gene correlations in the frontal cortex area (Figure 6 a-d). Prompted by these observations, we identified key players in the TCX AD network, including 46 protein-coding genes and 27 lncRNAs (Table 2 and 3). Utilizing the established functions of protein-coding genes as an anchor, we bridged lncRNAs to the enriched biological processes, demonstrating the utility of network-based approaches. We not only nominated potential key lncRNAs involved in AD pathology but also predicted their roles in biological processes under both AD and non-AD conditions.

Networks provide a powerful framework for studying the structure of complex biological systems, offering valuable insights into gene-gene interactions. However, network analysis poses challenges, such as managing noise signals and enhancing the reliability of correlations. To address these issues, we employed several strategies to mitigate biases and increase robustness (Figure 1). First, we ensured that all datasets used had a minimum of 30 replicates per condition (Figure 2c), as larger sample sizes improve the accuracy of network inference (51,52). Second, we applied the wTO method (53) to compute gene-gene correlations. This approach accounts for shared network neighborhoods, producing more robust scores compared to simple correlation calculations, and incorporates both positive and negative correlations (54). The gene co-expression networks were subsequently filtered based on the absolute value of gene pair correlations. Following this, we integrated the networks within each brain region (TCX and FCX) (Figure 2b, c). This integration allowed us to treat them as biological replicates, identifying common interactions across the integrated networks and reducing the impacts of sampling and batch variability, a widely adopted approach in gene co-expression studies (37–40,55,56). Ultimately, this consensus network offers a high-resolution representation of the conserved cellular processes functioning under AD and control conditions, providing a robust foundation for further investigation.

Co-expressed genes are often coregulated or involved in similar pathways, forming densely interconnected clusters within the network. In this study, we identified gene clusters and their functions for each consensus network. Through a comprehensive comparative analysis of gene correlations and network topology between AD and control consensus networks, we observed remarkable network alterations. Specifically, in the TCX networks, substantial rewiring was evident within the AD clusters, which was also reflected in the links between protein-coding genes associated with enriched terms (Figure 4c, d). The emergence of new core genes (Figure 3a), along with their higher engagement in the AD network compared to the control network (18% versus 2%, Figure 4e), indicated a reorganization of the molecular landscape in AD, suggesting that specific genes adopted new roles within the altered regulatory environment. As a result, we identified 46 core protein-coding genes as candidates potentially playing a central role in the rewiring observed in enriched terms in the TCX AD network (Table 2).

Several of these genes have been supported by prior experimental or network-based studies as being linked to AD. Notably, 15 of these genes have been reported as hub genes in AD, including *ATL1* (57), *AP3M2* (58)*, ATP6AP2* (59), *ATP6V1A* (60), *ATP6V1D* (61,62), *ACSL3* (58,63), *CCNB1* (64), *GHITM* (62), *GRIA2* (65), *MDH1* (66), *MRPS30* (67), *MRPS35* (66), *NAPB* (68,69), *NDUFB5* (66) and *PFN2* (70). Several genes showed strong evidence supporting their role in AD pathology. For example, *REPL* knockdown led to increased Aβ levels and tau phosphorylation (71–73), *NAPB* was linked to neuroinflammatory neuronal damage (74), *MDH1* correlated with total and phosphorylated tau levels (74,75) and *VPS35* deficiencies or mutations caused altered endosomal trafficking and pathological tau accumulation (13,76–78). Of particular interest, *ABL1* (c-Abl) contributed to tau pathology through regulation of cytoskeletal signaling cascades (79,80). Our findings identified *ABL1* as an AD-specific hub annotated to the stress fiber assembly term (Table 2). Notably, about half of the 46 AD key genes were annotated to the mitochondrial energy production functional category. Among them, *NDUFA5* and *NDUFB5*, encoding subunits of Complex I of the mitochondrial respiratory chain, were consistently downregulated across TCX datasets (Figure 3c), aligning with their known significant downregulation in AD (81). Recent findings revealed that the endolysosomal V-ATPase acts as a receptor for Aβ and Tau in AD (14). Among the 13 genes involved in the V-ATPase complex, four—*ATP6V1B2*, *ATP6V1A*, *ATP6AP2*, and *ATP6V1D*—were in our key gene list. In addition to previously known AD- associated genes, several key genes identified in this study have been pursued as promising therapeutic targets for AD, including *ABL1* (82), *ACSL3* (63), *ATP6V1A* (60,83), *CISD* (84) and *COQ10B* (85–88).

Having identified many previously known AD-associated protein-coding genes with our network analysis, we leveraged the connectivity of lncRNAs with their neighboring protein-coding genes within the network to infer potential functions for these lncRNAs. Through this approach, 54 lncRNAs in the TCX AD network and 103 in the FCX control network were assigned to functional categories identified within each cluster (supplementary Table S8 and S11). The assigned biological functions were cross-validated by assessing the similarity of lncRNAs assigned to common functional categories between the TCX and FCX control networks, revealing significant overlap (Table 4). From the TCX AD network, 27 core lncRNAs with functional assignments were selected for further investigation (Table 3). We observed that all but six of these 27 lncRNAs were AD-specific core genes. Notably, *FBXW7-AS1*, *Lnc-PDE4D-2*, *Lnc-KLHL11-1*, *Lnc-NIF3L1-5*, *Lnc-KALRN-1*, *Lnc-CCDC68-1*, *HSALNG0099005*, *HSALNG0105911*, *ZRANB2-DT*, *ENSG00000287022*, and *SLC8A1-AS1* were functionally assigned exclusively within the AD network, suggesting a gain of function for these lncRNAs in the context of AD. Among them, *ZRANB2-DT*, assigned to neurogenesis and synapse signaling, stood out. This lncRNA acts as an antisense RNA for *ZRANB2* (Zinc Finger RANBP2-Type Containing 2), a gene previously reported to be associated with neuronal responses to chronic oxidative stress—a process closely linked to neurodegenerative diseases (89,90). We observed that the correlation between *ZRANB2-DT* and *ZRANB2* was AD-specific, suggesting a gain regulatory role for *ZRANB2-DT* in the context of AD. Additionally, differential gene expression analysis revealed that five of the candidate lncRNAs—*ITFG1-AS1*, *Lnc-MYCN-6*, *Lnc-EFR3A-6*, *HSALNG0089044*, and *Lnc-CCDC68-1*—were consistently dysregulated in AD across TCX datasets (Figure 3f, Table 3). Despite comprehensive data collection from databases such as MalaCards (91), LncRNADisease v3.0 (92), and existing literature (supplementary Table S13), none of the previously reported AD-associated lncRNAs overlapped with our candidate lncRNAs. Nevertheless, the presence of supportive evidence linking the candidate protein-coding genes to AD reinforced our confidence that some of the identified key lncRNA genes were also likely to play a role in AD, introducing promising new lncRNA candidates for further investigation and experimental validation.

Aligning with the current knowledge that AD-related pathology begins in the medial temporal lobe (2) and progresses to the frontal lobes in later stages (4), we had hypothesized that AD- related network disruptions would differ across brain regions. In the TCX AD network, we observed significant rewiring, while in the FCX AD network, there was a substantial loss of healthy gene correlations. This suggests that AD-related damage initially disrupts cellular interactions in the temporal lobe, as evidenced by the loss of control-specific links in the TCX control network. As the disease progresses, cells in the temporal lobe establish new gene expression patterns to compensate for the damage (AD-specific links). Conversely, the FCX AD network shows a predominant loss of gene correlations, suggesting that the frontal lobe may experience more extensive disruption before compensatory rewiring occurs. In line with this, studies have shown that genes like *ACSL3* (63) and those involved in mitochondrial respiratory complexes (81,93) display opposing patterns of dysregulation during the early and late stages of AD, potentially reflecting mechanisms of gene expression compensation as the disease progresses. This notion is further supported by findings from a single-cell study across 19 cortical regions, which identified the temporal lobe as exhibiting the earliest and most pronounced gene expression abnormalities (94).

While our study provided valuable insights into AD pathology through gene co-expression network analysis, it is important to acknowledge certain limitations inherent to the methodology. For instance, the thresholds applied for filtering links and annotating lncRNAs to biological processes may have introduced some biases. The threshold for filtering links may have removed interactions from one network if they fell below the chosen cutoff, leading to their classification as lost links in that network. However, we reasoned that if a link was weaker in one condition and stronger in another, it could still be considered an affected or altered correlation. Similarly, the strict threshold for annotating lncRNAs may have excluded some lncRNAs, as genes belonging to the same GO terms produced a wide range of RWR scores. While this strict threshold helped maintain confidence in the assignments, we recognized that non-annotated lncRNAs might still have been closely related to their neighboring protein-coding genes. Additionally, other genes within the network might have been crucial to AD pathology, but limitations in our methods may have prevented their identification. To address this latter limitation, we made the TCX AD network publicly available as an interactive Shiny app, enabling further exploration (see Data Availability). The app allows users to investigate the entire TCX AD network, including all clusters, and search for specific genes of interest. It also provides detailed information, such as gene neighbors, functional categories, differential expression status, core membership, and more, offering a resource for further discoveries and validations.

## Conclusion

In summary, our network-based approach allowed us to provide insights into the alterations of gene co-expression networks in AD brains, revealing clusters of functionally related genes driven by key protein-coding and lncRNA genes. While early stages of AD, particularly in the FCX, appear to be characterized primarily by a loss of gene interactions, disease progression seems to trigger network rewiring, potentially aimed at preserving fundamental brain functions. Additionally, we predicted functional roles for numerous lncRNAs within these networks, identifying 27 lncRNAs with potential involvement in AD pathology. These findings highlight promising candidates for future studies, offering deeper insights into their roles in both normal brain function and AD development.

## Material and Methods

### RNA-seq datasets

We downloaded the FASTQ files and metadata from the Synapse platform (95). A total of five RNA-seq datasets from two independent studies were used. The first study, known as the Mayo RNA-seq data (42) includes RNA-seq data for the temporal cortex human brain region. The FASTQ files for this study were downloaded from the Synapse repository with the Synapse ID syn3163039 (96). The second dataset, known as the Mount Sinai Brain Bank (MSBB) study (43), includes gene expression profiles for four brain regions: Brodmann area (BM) 36, BM44, BM22, and BM10. This dataset was downloaded from the Synapse repository with the Synapse ID syn3159438 (97). All samples were collected under precise clinical-pathological diagnoses and distinctive histologies, including Braak NFT stage (44), plaque density, CERAD (Consortium to Establish a Registry for Alzheimer’s Disease) (45), and CDR (clinical dementia rating) (46). Each AD and the corresponding control dataset from both brain areas consisted of an equal number of individuals across sexes with identical gene sets. Furthermore, these datasets have been carefully balanced for age, disease stage, and other potential confounders. All data underwent standardized preprocessing steps, including filtering based on library size and gene expression levels, to ensure uniformity across the datasets.

### Preprocessing of RNA-seq data

To ensure uniform data processing, we employed a unified pipeline that included standard alignment, quality control, and gene expression analyses on each dataset (Mayo and MSBB RNA-seq). The preprocessing steps included FASTQ quality control, adapter trimming, base quality control, and filtering out short reads. We excluded all FASTQ files with fewer than 12 million reads. Following this, reads were mapped to the human reference genome (hg38) using STAR (v 2.7.10b) (98). Samples with less than 60% unique mapped reads were also excluded from further analysis. We used the `featureCounts()`function from the Subread software (V 1.6.3) (99) for gene quantification, with the following parameters: `-s 0 -t gene -g gene_id -M -O --fraction`. `-s 0` (un-stranded data), `-t gene` (gene feature type), `-g gene_id` (grouped by gene ID), `-M` (allow multi-mapping reads), `-O` (count overlapping reads), and `--fraction` (fractional assignment of multi-mapping reads). To account for both sense and antisense lncRNAs, reads were mapped specifically to the gene feature type (`-t gene`) and counted fractionally by `countMultiMappingReads` arguments (`--fraction`). Noteworthy, the human gene annotation file version 103 (*Homo_sapiens.GRCh38.103.gtf*) from Ensembl was used for gene annotation. Notably, both datasets contained un-stranded sequence data, which required the use of the `-s 0` option.

To remove systematic effects in the RNA-seq data, we normalized all gene count data using the `DESeq2` R package (100), which normalizes for both sequencing depth and RNA composition. Moreover, genes with low expression levels can introduce noise into the correlation calculation process. Hence, we excluded genes with low expression levels to enhance the reliability of the correlation analysis. The criteria for excluding low-expressed genes were set such that only genes with a count of at least 1 count per million (cpm) in 70% of the samples were retained. The use of cpm avoids favoring genes expressed in larger libraries.

### Signed weighted co-expression network

Gene co-expression networks represent gene-gene interactions, with nodes representing genes and edges indicating the strength of their co-expression (101). The weight of an edge reflects the strength of the co-expression relationship between two genes, while the sign distinguishes between positive and negative correlations, showing whether genes have similar or opposing expression patterns. The signed weighted gene co-expression networks were constructed using the weighted Topological Overlap (wTO) method separately for each RNA-seq dataset. Using the wTO method, link weights for connected nodes are calculated through an averaging process that accounts for all their common connections (47). The wTO calculation was performed using the `wTO` R package. In summary, for each dataset, Pearson’s correlation coefficient (ρ) was calculated for all pairs of genes to generate the adjacency matrix A = [a_i,j_], where each element a_i,j_ represents the correlation between gene i and gene j. The weighted Topological Overlap (ω_i,j_) was then calculated for each pair of nodes i and j, reflecting the shared network connectivity between the two genes. Gene co-expression correlations with |*ω_i,j_*| < 0.5 were discarded. This threshold was chosen to balance the inclusion of meaningful interactions while reducing the influence of weak or spurious associations. This threshold, higher than the third quartile value (mean of 0.4) and close to the maximum observed weight (mean of 0.61), ensures that only the strongest links are preserved for further analysis (Figure 2e, supplementary Table S1).

### Consensus network construction by integrating wTO networks

To obtain a single consensus network derived from multiple independent wTO networks, the `intersection()`function from the `igraph` package was used. A custom function was developed to subset the overlapping links across the given networks and calculate an average weight for each link. Links with differing signs were excluded. The consensus weight between nodes i and j (*wTO_CN_*= [*Ω_i,j_*]) for common links across n replicate networks (`k=1,…,n`) was calculated using the formula from the `wTO.Consensus()` function of the `wTO` R package, as follows:

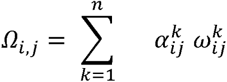

Where

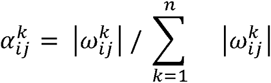

This approach ensures that stronger link weights 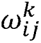 contribute more significantly to the corresponding consensus weights through a greater weighting factor 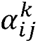. This helps preserve the impact of strong links, preventing their dilution by weaker ones. In contrast, a simple average would treat all connections equally, potentially blurring the distinction between strong and weak links and resulting in the loss of valuable information.

### Differential co-expression networks analysis

To identify differences between AD and control consensus networks, we performed a network comparison analysis using the `CoDiNA` R package (49). CoDiNA (Co-Expression Differential Network Analysis) is a network comparison framework that systematically identifies shared and condition-specific gene-gene interactions between multiple networks. Instead of simply highlighting differentially connected genes, CoDiNA classifies links into three categories: (1) conserved links that are common between two conditions with the same sign (α-link); (2) condition-specific links, appearing only in one condition (β-link); and (3) reversed links, representing common correlations with opposite signs in the two conditions (γ-links). This qualitative framework enables a clear interpretation of how gene interactions differ between conditions. For example, conserved links indicate stability in gene regulation, while condition-specific and reversed links highlight shifts in regulatory relationships that may underpin disease pathology. The package allows comparison of multiple networks, requiring input in a table format where the first two columns specify node pairs, and the third column contains their correlation values. We disabled weight normalization (stretch = FALSE) and applied the default cutoff (0.33) to define the minimum weight threshold for a link to be considered present. CoDiNA considers genes (nodes) present in both networks to avoid false associations, as the absence of a gene in one network might be due to technical reasons. However, given the nature of the gene expression data used in this study to construct the wTO consensus networks, the gene lists were identical for both conditions. Thus, the absence of a gene in one network did not necessarily indicate a technical issue but rather that it might not meet the criteria for meaningful correlations, such as the weight cutoff and network integration. To address this, we added the absent genes to each network with a zero-weight assigned to their links, ensuring that genes present in one condition but absent in the other yet still exhibiting a correlation were not excluded from the comparison. This approach enabled a comprehensive analysis of network differences between AD and control conditions.

### Hub core genes

To determine hub genes within each network, we used the K-core decomposition method (102), which identifies the most interconnected nodes by progressively removing nodes with degrees less than k until no more such nodes exist. K-cores are determined as subgraphs where every vertex has a degree of at least k. Therefore, vertices with the highest coreness scores represent the most tightly integrated and connected parts of the network. The k-core subgraph is defined through the following process: first, all nodes and their links are removed if their degree is less than k (starting with). The algorithm works iteratively, increasing k each time, until no more nodes meet the criteria (Figure 7). This effectively peels off layers (shells) of nodes until the graph has none left. For each network, all nodes were classified into shells based on the coreness scores, and nodes with the highest score were assigned as core hub genes of that network. We used the `coreness()` function from the `igraph` R package to calculate the coreness scores.

**Figure 7:**
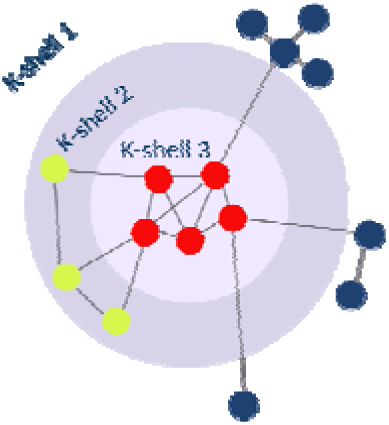
Illustration of K-core decomposition method. (103). All nodes of the network are classified into shells based on their coreness score. The inner shell (K-shell 3) with red-colored nodes represents those with the highest connectivity within the network, considered as core hub nodes in this study.

### Identifying sub-network clusters

Clusters are groups of genes with similar expression profiles and were explored based on the topological structure of the gene co-expression network. The Louvain method (104) was used for clustering. While this function can consider edge weights, it only accepts positive numeric vectors for weights. Since the networks contain both positive and negative weights, the weights argument was set to NULL, ensuring that all links are considered without their signs. Additionally, the resolution argument was set to 1, as it corresponds to the original definition of modularity defined by the algorithm. Essentially, a lower resolution value results in fewer and larger clusters. Cluster smaller than 20 links were excluded from further analysis.

### GO enrichment analysis

Gene Ontology (GO) enrichment analysis for biological process terms was performed using the `compareCluster()`function from the `clusterProfiler` R package (105). The analysis was conducted separately for each gene cluster, using the same background genes. Background genes comprised the protein-coding genes that were included in the wTO construction. P-values were adjusted using the false discovery rate (FDR) method, with a significance threshold of 0.05.

### Redundancy reduction of the enriched GO terms

Although GO enrichment is a powerful tool for understanding the biological processes and pathways associated with a set of genes, this analysis is frequently challenged by the redundancy inherent in GO terms. This redundancy occurs because of the hierarchical nature of the GO hierarchy, where parent terms frequently encompass child terms. For example, a gene may be annotated with both a parent term and a more specific term at different levels of the hierarchy within the GO. Various methods have been developed to address this issue, each with its own advantages and disadvantages. However, none of these methods work flawlessly, and some incorrectly group GO terms with inappropriate representative parent terms. Therefore, a manual approach was chosen to mitigate redundancy. Child terms were manually grouped into broader cellular biological process terms to which they belong. Additionally, GO terms that were irrelevant, included fewer than five genes, or represented different terms for the same biological process were excluded.

### Random Walk with Restart

The random walk on a graph is an iterative process where a walker transitions from its current node to a randomly selected neighbor, starting from specified source node(s), known as seeds. The Random Walk with Restart (RWR) algorithm extends this concept by allowing the walker to restart at the seed node(s) with a probability r at each time step (106). In this study, RWR was used to assign lncRNAs within each cluster to the enriched functional categories (GO terms) identified for that cluster. Each co-expression network constructed in this study included both protein-coding genes and lncRNAs as nodes. Knowing the involvement of protein-coding genes in the enriched biological processes, we aimed to assign lncRNAs to these processes by leveraging their connectivity with neighboring protein-coding genes within the network. For each cluster, we performed iterative RWR calculations, using genes from each functional category as seed genes and ranking lncRNAs based on their associations. The process for each iteration included the following steps: (1) performing RWR calculations within each network cluster, (2) using genes from each functional category within the cluster as seed genes and computing RWR scores for lncRNAs in that cluster, (3) excluding link weights when computing RWR scores due to the algorithm’s inability to handle negative values in signed networks, (4) computing the adjacency matrix for the network, (5) normalizing the adjacency matrix by columns, and (6) setting the restart probability (`r`) to 0.7.

### Assigning lncRNAs to the functional category based on RWR scores

LncRNAs were ranked based on their computed scores for each functional category, and those scoring above the 10th percentile were assigned to that functional category. As cross-validation, we calculated RWR scores for genes within each functional category using a leave-one-out method. Within each cluster, for each protein-coding gene in a given GO group, we treated all other genes except one as seeds and computed the RWR score for the excluded gene. This process was repeated across all genes annotated to each functional category. The assigned lncRNA was accepted only if its RWR score did not fall below the lowest RWR scores observed in the leave-one-out calculation for that category (Figure S1).

### Statistical evaluation tests

Szymkiewicz-Simpson coefficient and permutation test was specifically chosen to rigorously assess whether the observed overlaps in lncRNA annotations were significant or merely due to random chance. The similarity between assigned lncRNAs to a common functional category across the two networks was initially measured using the Szymkiewicz-Simpson coefficient. For each functional category, the overlap coefficient was calculated by determining the intersection of lncRNAs assigned to the respective functional category in each network, normalized by the minimum number of lncRNAs assigned to that functional category in either network. This approach normalized the overlap relative to the size of the smaller set, accommodating the significant disparity in the total number of lncRNAs between the FCX and TCX control networks (1100 and 504, respectively). The statistical significance of the observed overlap was assessed using a permutation test. The permutation test involved generating 10,000 random permutations of lncRNA sub-samples from the complete list of lncRNAs for each network. During each permutation, lncRNAs were randomly sampled from the entire pool of lncRNAs in each network to create a new, synthetic group of lncRNAs matching the original count of lncRNAs assigned to that functional category. The significance of the observed overlap was assessed by calculating the proportion of permuted overlap coefficients that were equal to or greater than the observed coefficient, which was then divided by the number of permutations to yield a p-value.

To assess the difference in lncRNA representation between TCX AD and control core networks, we constructed a contingency table comparing the number of core lncRNAs in each network and evaluated statistical significance using Fisher’s exact test.

### Differential gene expression analysis

The `DESeq2` R package was used to identify differentially expressed genes (DEGs) in AD compared to control. Genes with a false discovery rate (FDR)LJ<LJ0.05 and absolute log fold changeLJ> 0.3 were considered significant DEGs. Due to the integration of several networks to make each consensus network, identifying DEGs presented a challenge. To address this, only DEGs that were consistently identified across all gene expression datasets used to construct the given network were selected. For example, the AD TCX consensus network integrates three networks: AD Mayo, AD MSBB BM36, and AD MSBB BM22. DEGs were identified separately for each dataset—AD Mayo, AD MSBB BM36, and AD MSBB BM22. Ultimately, the DEGs in the AD TCX consensus network were determined by selecting those DEGs that overlapped across all these three datasets.

### Network visualization

The network visualizations presented in the manuscript were created using Cytoscape v.310.2 (107) and Gephi v.0.10.1 (48).

### Single cell RNA-seq analysis

The single-cell RNA sequencing data was downloaded from the Synapse repository with the Synapse ID syn22079621. This dataset includes samples from the prefrontal cortex, comprising 7 control and 11 AD cases (108). Data preprocessing and analysis were performed using Seurat v 4.3.0.1 (109). To ensure high-quality single-cell data, we applied the following filtering criteria: cells with fewer than 200 detected genes or more than 7500 detected genes were removed, and cells with more than 5% of reads mapping to mitochondrial genes were excluded to eliminate low-quality or stressed cells. Following quality control, the dataset was normalized using the `NormalizeData` function in Seurat. Uniform Manifold Approximation and Projection (UMAP) was used for dimensionality reduction and visualization of cellular clusters. Clusters were identified and cell type were manually annotated based on established marker genes including *AQP4*, *SLC1A2*, and *GPC5* for astrocytes; *MBP*, *MOBP*, and *PLP1* for oligodendrocytes; *CD74*, *LPAR6*, *DOCK8*, and CSF1R for microglia; FLT1, *CLDN5*, and *LAMA2* for endothelial cells; *VCAN*, *DSCAM*, *PCDH15*, and *MEGF11* for oligodendrocyte progenitor cells; *NRGN* for excitatory neurons; and *GAD1* for inhibitory neurons.

### Bulk RNA-seq deconvolution

To estimate cell-type proportions in bulk RNA-seq data, deconvolution was performed using `CIBERSORT` R package (110). The single-cell dataset described earlier was used to construct a signature gene matrix, utilizing only control samples for this purpose. The expression data was normalized using `SCTransform` (SCT), and the signature matrix was generated from SCT-normalized data by averaging expression across cell types. The CIBERSORT algorithm was run with 100 permutations, and only samples with a p-value < 0.05 were retained for further analysis. The estimated cell-type proportions between AD and control samples were compared using the Wilcoxon test, with false discovery rate-adjusted p-values < 0.05 considered significant.

### Data availability

The datasets used in this study are available in the Synapse online repositories under the following accession IDs: Mayo RNA-seq dataset (Synapse ID: syn3163039) (96) and MSBB RNA-seq dataset (Synapse ID: syn3159438) (97).

### R script code availability

To enhance the exploration of our findings, we provided an interactive Shiny app that visualizes the gene co-expression networks. This tool enables researchers to investigate specific genes and their correlations and associated functions, offering deeper insights into the TCX AD and control networks. They can be downloaded from https://github.com/mpstar92/Shiny_App_ADnet.

The R script code used in this study is available on the GitHub repository at https://github.com/Tima-Ze/AD_network.

## Supporting information

Suppemental material

## Acknowledgments

Data used in this project were obtained from the Synapse platform (https://www.synapse.org/), including the Mayo RNA-seq study (96), the Mount Sinai Brain Bank study (MSBB) dataset (97) and Human AD Single-nuclei Multi-Omics (108). We extend our gratitude to the data providers on the Synapse platform, as well as the researchers and authors involved in these studies. This study was supported by the International Max Planck Research School for Biology and Computation (IMPRS-BAC), Berlin, Germany. We would like to thank Vladimir M. Jovanovic, Melanie Sarfert, Yao-Chung Chen, and Rebecca Sinead Saager for their kind assistance and contributions, as well as Ralf Herwig for his scientific methodology advice.

## Author contributions

Nowick supervised the research. Zebardast analyzed and visualized the data and wrote the manuscript. Riethmüller developed the Shiny app. All authors read and approved the final manuscript.

